# Individual-subject functional localization increases univariate activation but not multivariate pattern discriminability in the ‘multiple-demand’ frontoparietal network

**DOI:** 10.1101/661934

**Authors:** Sneha Shashidhara, Floortje S. Spronkers, Yaara Erez

## Abstract

The frontoparietal ‘multiple-demand’ (MD) control network plays a key role in goal-directed behavior. Recent developments of multivoxel pattern analysis (MVPA) for fMRI data allow for more fine-grained investigations into the functionality and properties of brain systems. In particular, MVPA in the MD network was used to gain better understanding of control processes such as attentional effects, adaptive coding, and representation of multiple task-relevant features, but overall low decoding levels have limited its use for this network. A common practice of applying MVPA is by investigating pattern discriminability within a region-of-interest (ROI) using a template mask, thus ensuring that the same brain areas are studied in all participants. This approach offers high sensitivity, but does not take into account differences between individuals in the spatial organization of brain regions. An alternative approach uses independent localizer data for each subject to select the most responsive voxels and define individual ROIs within the boundaries of a group template. Such an approach allows for a refined and targeted localization based on the unique pattern of activity of individual subjects while ensuring that functionally similar brain regions are studied for all subjects. In the current study we tested whether using individual ROIs leads to changes in decodability of task-related neural representations as well as univariate activity across the MD network compared to when using a group template. We used three localizer tasks to separately define subject-specific ROIs: spatial working memory, verbal working memory, and a Stroop task. We then systematically assessed univariate and multivariate results in a separate rule-based criterion task. All the localizer tasks robustly recruited the MD network and evoked highly reliable activity patterns in individual subjects. Consistent with previous studies, we found a clear benefit of the subject-specific ROIs for univariate results from the criterion task, with increased activity in the individual ROIs based on the localizers’ data, compared to the activity observed when using the group template. In contrast, there was no benefit of the subject-specific ROIs for the multivariate results in the form of increased discriminability, as well as no cost of reduced discriminability. Both univariate and multivariate results were similar in the subject-specific ROIs defined by each of the three localizers. Our results provide important empirical evidence for researchers in the field of cognitive control for the use of individual ROIs in the frontoparietal network for both univariate and multivariate analysis of fMRI data, and serve as another step towards standardization and increased comparability across studies.

## 1. Introduction

Multiple studies have provided consistent evidence for the involvement of a large distributed network of frontal and parietal regions in cognitive control and flexible goal-directed behavior (Desimone & Duncan, 1995; Duncan, 2006, 2010, 2013; Duncan & Owen, 2000; Fedorenko, Duncan, & Kanwisher, 2013; Stiers, Mennes, & Sunaert, 2010). This network has been termed the ‘multiple-demand’ (MD) network (Duncan, 2006), and it closely resembles other networks that have been associated with control processes such as the cognitive control network (e.g. Cole and Schneider, 2007), task-activation ensemble (Seeley et al., 2007), and task-positive network (Fox et al., 2005). The MD network includes the intraparietal sulcus (IPS), the anterior-posterior axis of the middle frontal gyrus (MFG), the anterior insula and adjacent frontal operculum (AI/FO), the pre-supplementary motor area (pre-SMA) and the dorsal anterior cingulate (ACC) (Duncan, 2010, 2013). A primary characteristic of this network is an increase in activity with increased demand, especially seen through functional magnetic resonance imaging (fMRI) blood oxygenation level dependent (BOLD) activity, across a variety of cognitive domains such as working memory, task switching, inhibition, math, and problem solving (Cole & Schneider, 2007; Dove, Pollmann, Schubert, Wiggins, & Yves von Cramon, 2000; Fedorenko et al., 2013; Shashidhara, Mitchell, Erez, & Duncan, 2019).

The development of multivoxel pattern analysis (MVPA) methods (Haxby et al., 2001; Haynes & Rees, 2006; Kriegeskorte, Goebel, & Bandettini, 2006) has further led to a variety of findings related to the representation of multiple aspects of cognitive control across the MD network. These include attentional effects, adaptive coding, and coding of target features and task rules (Erez & Duncan, 2015; Etzel, Cole, Zacks, Kay, & Braver, 2016; Nee & Brown, 2012; Nelissen, Stokes, Nobre, & Rushworth, 2013; Wisniewski, Goschke, & Haynes, 2016; Woolgar, Thompson, Bor, & Duncan, 2011). MVPA allows for a fine-grained investigation of distributed patterns of activity and the information that is conveyed in these patterns related to different experimental conditions and their respective cognitive constructs. However, its use in the frontoparietal MD network has been limited by overall low decoding levels, or discriminability between conditions, compared to other brain systems (Bhandari, Gagne, & Badre, 2018). This could be, at least in part, due to differences between individuals in respect to the spatial organization of the network. Indeed, it has been demonstrated that across the fronto-parietal lobes, several close by regions have been fractionated into different networks in resting state studies, with different boundaries in individuals (Glasser et al., 2016; Schaefer et al., 2018; Yeo et al., 2011). Nevertheless, many MVPA studies of the MD network have used a group template to define the same regions of interest (ROIs) for all subjects and to investigate task-related representations. Such a group template is robust and easily comparable across studies. However, it provides high sensitivity at the cost of specificity to individual differences, as it might not accurately identify regions in individual subjects due to both anatomical and functional differences (Brett, Johnsrude, & Owen, 2002; Fedorenko et al., 2013; Nieto-Castañón & Fedorenko, 2012).

An alternative to the group template is using an independent functional localizer task to establish subject-specific ROIs. In this approach, participants perform a short task in addition to the main task in the scanning session, and the data from this task is used to localize regions-of-interest to be tested with data from the main task. This method is commonly used in vision research (Erez & Yovel, 2014; Lafer-Sousa, Conway, & Kanwisher, 2016; Reddy & Kanwisher, 2007; Weiner et al., 2018). For example, specific tasks are used to identify regions in individuals that are recruited for face processing (Berman et al., 2010; Kanwisher, McDermott, & Chun, 1997) and object processing (Malach et al., 1995). Task contrasts such as faces versus scrambled faces or objects versus scrambled objects are applied, and the experimenter can identify ROIs as clusters of activity in individual subjects. However, this alternative of manually defining ROIs by the experimenter using a functional localizer is subjective and therefore may be prone to biases, inaccuracies, and reduced reproducibility (Garrison et al., 2015; Krishnan, Slavin, Tran, Doraiswamy, & Petrella, 2006).

To overcome the limitations of both the group template and the individual manually-defined regions, a hybrid group-constrained subject-specific approach has been proposed for use in the language system (Fedorenko, Hsieh, Nieto-Castañón, Whitfield-Gabrieli, & Kanwisher, 2010; Fedorenko, Nieto-Castañon, & Kanwisher, 2012), and later expanded to the ventral pathway of the visual system (Julian, Fedorenko, Webster, & Kanwisher, 2012) and theory of mind regions (Paunov, Blank, & Fedorenko, 2019). In this approach, independent subject-specific localizer data are collected in addition to the main experimental task, then the thresholded contrast data from this task are masked with a group template of regions and only the voxels that were responsive to the localizer task within the group template are used for further analysis in the main experimental task. The advantage of this approach is the use of a group template that ensures targeting of similar areas for all participants, as well as refining this localization by subject-specific activations within these areas. It therefore offers an objective experimenter-independent definition of subject-specific regions that does not require manual region definition. Importantly, it supports comparability across different studies because selected voxels in individual subjects are constrained to a group template. Using this group-constrained subject-specific ROIs approach has been shown to increase the detected univariate BOLD response associated with contrasts of interest in language-related areas compared to when a group template was used as ROIs (Fedorenko et al., 2010). The benefit of using individually defined ROIs for univariate results has also been shown for the visual system (Saxe, Brett, & Kanwisher, 2006) and has been modeled and demonstrated using simulation data (Nieto-Castañón & Fedorenko, 2012).

The hybrid group-constrained subject-specific approach has been subsequently used for the MD network in studies that used both univariate (Blank & Fedorenko, 2017; Blank, Kanwisher, & Fedorenko, 2014; Mineroff, Blank, Mahowald, & Fedorenko, 2018; Paunov et al., 2019) and multivariate measures (Erez & Duncan, 2015; Shashidhara & Erez, 2019) related to control processes. A spatial working memory task that has been previously demonstrated to robustly recruit the MD network was used as a localizer. In this localizer task, a highly demanding condition is contrasted with an easier version of the same task to identify the network in individual subjects, then constrained by an anatomical or group-average functional template to define the subject-specific ROIs. However, it remains an open question whether using these refined ROIs at the single subject level has any benefit for multivariate results in the MD network, similarly to the benefits that were previously reported for univariate results (Fedorenko et al., 2010; Nieto-Castañón & Fedorenko, 2012; Saxe et al., 2006). Using the MD group template for MVPA means that many voxels outside the individually-defined functional MD regions are included in the analysis, which may not express the domain-general characteristics of the MD network. In fact, they may be part of other nearby brain systems (Glasser et al., 2016; Schaefer et al., 2018; Yeo et al., 2011), and their inclusion in the multivariate analysis may potentially mask out pattern-based differences between the experimental conditions of interest. If this is indeed the case, then the identification of the MD network in individual subjects has the potential to critically improve our ability to detect the neural signature of control processes as measured by multivariate methods, thus substantially increasing the benefit of using MVPA in cognitive control research. On the other hand, it has been previously demonstrated in the visual system and using simulations that even voxels outside the regions of increased univariate activity contribute to multivoxel discrimination (Haxby et al., 2001; Kriegeskorte et al., 2006). This implies that using more finely defined subject-specific ROIs instead of the large group template may potentially reduce discrimination levels. A related point concerns the size of ROIs. Increased decoding levels have been previously linked to increased size of ROI, at least in the visual system (Eger, Ashburner, Haynes, Dolan, & Rees, 2008; Said, Moore, Engell, Todorov, & Haxby, 2010; Walther, Caddigan, Fei-Fei, & Beck, 2009), highlighting the importance of controlling ROI size for MVPA. Using the group-constrained subject-specific ROIs allows for such control by selecting a fixed number of voxels from each ROI with the largest localizer contrast values. Such control for ROI size enables the comparison of decoding levels between different regions within the network, which vary in size, as well as with regions outside this network. This, however, should not come at the expense of reduced decodability, if indeed the use of smaller ROIs reduces pattern discriminability. Overall, existing data provide only limited evidence regarding the link between the use of subject-specific ROIs and multivoxel pattern measures in the MD network.

In the current study, we build on the previously reported findings and ask whether using functionally defined subject-specific ROIs affects multivoxel pattern results in the MD network. Because the recruitment of the MD network at the group level is observed across a range of cognitive domains (Fedorenko et al., 2013), different tasks can be potentially used as localizers. We therefore also ask whether different localizers may have different effects on multivariate results. To address these questions, we use three localizer tasks and an independent rule-based criterion task. The localizer tasks are spatial working memory, verbal working memory, and a Stroop-like task, which have all been previously shown to consistently recruit MD regions (Fedorenko et al., 2013). We first assess the reliability and variability of the level of recruitment of the MD network by the localizers. We then assess the benefit of the subject-specific ROIs for univariate results in the independent criterion task, aiming to replicate and generalize previous findings. Finally, we systematically test for the effect of using the subject-specific ROIs defined by each of the three localizers on multivariate results in the independent criterion task. We provide important empirical evidence for the effect of using subject-specific ROIs on the ability to detect task-related representations in the MD network as reflected in distributed patterns of activity.

## 2. Methods

### 2.1. Participants

A total of 25 healthy participants (18 female, mean age 23.8 years) took part in the study. Three participants were excluded because of movements larger than 5 mm during at least one of the scanning runs, and one participant was excluded due to slice by slice variance larger than 300 after slice time correction in more than two runs. In addition, two participants were excluded due to technical problems with the task scripts. Lastly, one participant was excluded to maintain the balance of the order of localizers across participants. This participant was chosen randomly out of the participants who had the same order of localizers and prior to any data analysis beyond pre-processing. Overall, 18 participants were included in the analysis. All participants were right-handed and had normal or corrected-to-normal vision. Participants were either native English speakers or had learnt English at a young age and received their education in English. Participants gave written informed consent prior to participation and received a monetary reimbursement at the end of the experiment. Ethical approval was obtained from the Cambridge Psychology Research Ethics Committee.

### 2.2. Experimental paradigm

The study consisted of three localizer tasks and one rule-based similarity judgment task. The localizers were: a spatial working memory (WM) task, a verbal working memory task and a Stroop task, variations of which have previously been shown to recruit the MD network (Fedorenko et al., 2013). The rule-based task was used as a criterion task to test for univariate effects and rule decoding using MVPA, using both subject-specific ROIs based on activation data from the localizers and the group template. Participants practiced all tasks before the start of the scanning session. During scanning, participants performed two runs of each localizer followed by four runs of the rule-based task. The two runs of the same localizer always followed each other, and the order of the three localizers was balanced across participants. The average total scanning session duration was 105 minutes.

### 2.3. Localizer tasks and criterion task

The spatial WM and verbal WM localizer tasks were adapted from Fedorenko et al. (2013) and the Stroop task was adapted from Hampshire et al. (2012). The localizers were chosen based on their consistent recruitment of the MD network as has been shown by Fedorenko et al. (2013). The localizers all followed a blocked design. Each run contained 10 blocks, alternating between Easy (5 blocks) and Hard (5 blocks) task conditions. There were no indications for the start or end of each block. The localizer tasks were designed to be used with a contrast of Hard versus Easy conditions, and in order to keep them as short as possible they did not include fixation blocks. The first run always started with an Easy block, and the second with a Hard block. All blocks lasted for 32 seconds, leading to a total run duration of 5 minutes and 20 seconds.

All tasks were coded and presented using Psychtoolbox (Brainard, 1997) for MatLab (The MathWorks, Inc.). Stimuli were projected on a 1920 x 1080 screen inside the scanner, and participants used a button box, with one finger from each hand to respond.

#### 2.3.1. Localizer 1: spatial WM task

In the spatial WM task (**Figure 1A**), each trial started with an initial fixation dot (0.5 s), followed by a 3×4 grid with either one (Easy condition) or two (Hard condition) highlighted cells. The highlighted cells were displayed over four seconds, with different cells highlighted every one second, leading to an overall four (Easy condition) or eight (Hard condition) highlighted cells in each grid. In a subsequent two-forced choice display, two grids with highlighted cells were presented on the right and left sides of the screen. Participants pressed a button (left or right) to indicate which grid matched the previously highlighted cells. After a response window of 3.25 s, a feedback was presented for 0.25 s. The correct grid appeared an equal number of times on the right and left. Overall, each trial was 8 s long, and each task block contained four trials.

**Figure 1.**
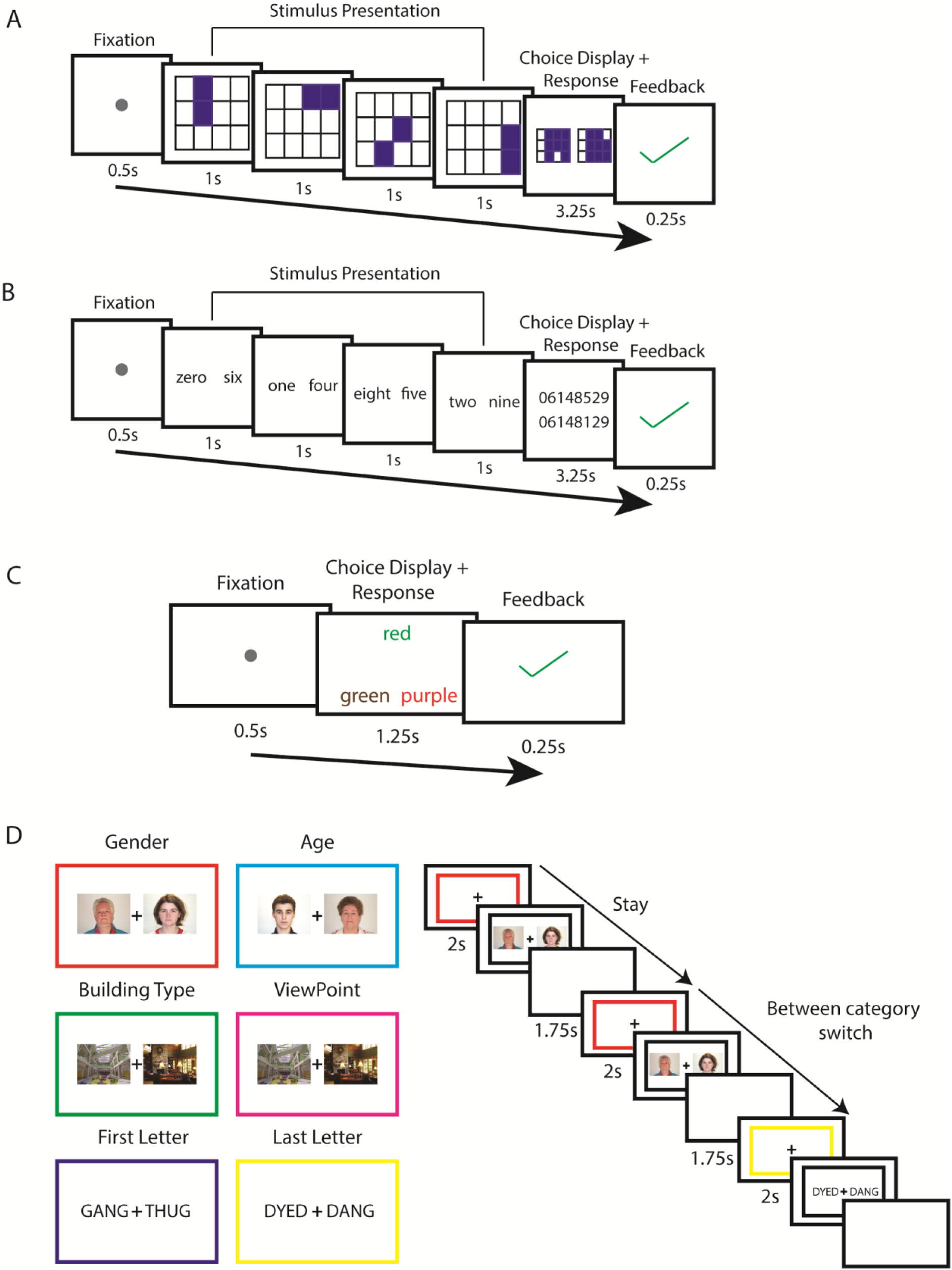
Schematic overview of the Hard condition of the three localizer tasks and the rule-based similarity judgment task. **A:** Spatial working memory task. Participants were presented with highlighted cells in a 3×4 grid, on four consecutive screens. In the Easy condition, one cell was highlighted at a time, and in the Hard condition two cells were highlighted in each screen. They selected the grid with the correctly highlighted cells in a subsequent two-forced choice display. They received feedback after each trial. Positive feedback was indicated by a green tick, and negative by a red cross. **B: Verbal working memory task.** A design similar to the spatial working memory task was used, with written digits instead of a grid. **C: Stroop task.** In each trial, three color names were presented. Participants selected the answer option (out of two options at the bottom) that described the ink color of the test word on top. In the Easy condition, the ink color of the test word matched its color name (congruent), and in the Hard condition they were different (incongruent). In this example of a Hard trial, the correct answer is the word ‘green’ written in brown, and the ink color of the distractor (the word ‘purple’ written in red ink) matches the color name of the test word, thus increasing the difficulty level. See the text (*2.3.3*) for more details. For all localizer tasks, feedback was presented at the end of each trial. **D: Rule-based similarity judgment task.** The six rules with the corresponding colored frames, paired by the category domain that they should be applied on (left). In each trial, a colored frame indicated the rule, followed by two images. Participants indicated whether the images are the ‘same’ or ‘different’ based on the rule. Transitions between trials could be either the same rule and category domain (‘Stay’), a switch of rule within the same category domain (‘within-category Switch’), or a switch of rule between category domains (‘between-category Switch’). In this example (right), participants indicated whether the two faces have the same gender or not in the first trial, followed by a Stay trial and a between-category Switch trial.

#### 2.3.2. Localizer 2: verbal WM task

The verbal WM task (**Figure 1B**) followed a similar design to the spatial WM task. Following fixation (0.5 s), participants were presented with four consecutive screens containing one (Easy condition) or two (Hard condition) written digits. In a following two choice display, participants indicated the correct sequence of digits by pressing a button. The two answer options were displayed at the center of the screen, one above the other for ease of reading. The left button was used to choose the sequence on top, while the right button was used for the sequence at the bottom. The correct sequence appeared an equal number of times on the top and bottom. Following a response window of 3.25 s, participants were given feedback at the end of each trial (0.25 s). Each trial was 8 s long, and each task block contained four trials.

#### 2.3.3. Localizer 3: Stroop task

The third localizer was a variation of the Stroop task (**Figure 1C**). On each trial, following a fixation dot (0.5 s), participants were presented with a test word, which was the name of a color, written in color at the top of the screen. In the Easy condition, the ink color was the same as the color name (congruent) (e.g. the word ‘green’ written in green ink), and in the Hard condition the ink color and the color name were different (incongruent) (e.g. the word ‘red’ written in green ink). Participants had to indicate the ink color (rather than the written color name) by choosing one of two answer options at the bottom of the screen, displayed at the same time. The answer options were color name words, and their ink color was different from its name. Participants had to choose the word, i.e. the written color name (regardless of the ink color) that matched the ink color of the test word at the top. Therefore, participants had to switch between attending to the ink color of the test word (ignoring the written color name) to detecting the matching written color name (and ignoring the ink color) out of the two answer options at the bottom. In the congruent condition, the ink color of the answer options was chosen randomly, excluding the color name (and ink color) of the test word (e.g. the options for the above congruent test word example could be the word ‘blue’ written in brown ink and ‘green’ written in purple ink, with the latter being the correct answer). In the incongruent trials, the ink color of one of the answer options matched the color name (and not the ink color) of the test word. On half of the incongruent trials, it was the correct answer that had the same ink color as the test word color name (for example, if the test word is ‘red’ written in green, then a correct answer could be ‘green’ written in red). On the other half, it was the incorrect option with ink color the same as the test word color name (e.g. if the test word is ‘red’ written in green, then the incorrect option could be ‘purple’ written in red, see example in Figure 1C). This was done to further increase the conflict between stimuli and thus the difficulty level of the hard blocks while ensuring that the ink color of the test word cannot be used when choosing an answer. A total of six colors were used in this task (red, green, blue, orange, purple, and brown). Participants had 1.25 s to view the stimuli and respond, after which they received feedback for 0.25 s. Each trial was 2 s long, and blocks consisted of 16 trials each.

#### 2.3.4. Criterion rule-based similarity judgment task

The rule-based similarity judgment task was a variation of a task previously used by Crittenden and colleagues (2016, 2015) and Smith et al. (2018). This was chosen as the criterion task because it allowed for testing both univariate and multivariate effects. Univariate effects were addressed using the task switching aspect of the task. Multivariate effects were addressed using rule decoding, which has been previously observed across the MD network using this task. Additionally, this task enabled a more detailed investigation of the potential effect of using individual ROIs on the difference between two types of rule discriminations.

Prior to the start of the scanning session, participants learned to associate colored frames with six rules (**Figure 1D**). In each trial, participants indicated whether two displayed images were the same or different based on the given rule. The six rules were applied on stimuli from three different category domains (faces, buildings, words), with two rules per category. The rules and category domains were: (1) Gender (male/female, red frame) and (2) Age (old/young, light blue frame) applied on faces; (3) Building type (cottage/skyscraper, green frame) and (4) Viewpoint (seen from the outside/inside, magenta frame) applied on buildings; (5) First letter (dark blue frame) and (6) Last letter (yellow frame) applied on words and pseudo-words.

Each trial began with a colored frame (2 s) that indicated the rule to be applied. This was followed by two stimuli presented to the left and right of a fixation cross (**Figure 1D**) and the cue replaced by a black frame. Participants had to respond ‘same’ or ‘different’ based on the rule by pressing the left or right button, and response mapping was counterbalanced across subjects. The task was self-paced and the stimuli were on the screen until a response was made. Each trial was followed by an inter-trial interval (ITI) of 1.75 s. The colored frame was 14.96° visual angle along the width and 11.60° along the height. The choice stimuli were displayed at 3.68° eccentricity from the center and were 6.0° and 4.5° along the width and height.

We used an event-related design, with 12 trials per rule in each run. Out of these, half (6) of the trials had ‘same’ as correct response and half (6 trials) had ‘different’ as correct response. Out of the 6 trials with ‘same’ as correct response, half (3 trials) also had ‘same’ as correct response if the other rule for the category was applied, therefore identical responses using either rule for this category. The other half (3 trials) had ‘different’ as correct response using the other rule in this category, therefore different responses for the two rules. A similar split was used for the 6 trials with ‘different’ as the correct response. To decorrelate the cue and stimulus presentation phases, 4 out of 12 trials of each rule were chosen randomly to be catch trials, in which the colored frame indicating the rule was shown, but was not followed by the stimuli.

The task included switches between rules that were used for the univariate analysis. Trial switches were defined based on the rules in two consecutive trials. In ‘Stay’ trials, the previous trial had the same rule and category domain. For ‘within-category Switch’ trials, the previous trial had the same category domain (faces, buildings, or letters) but a different rule (e.g. age vs. gender for faces). For ‘between-category Switch’ trials, the previous trial had a different category domain and therefore necessarily a different rule. There was an equal number of Stay, within-category Switch and between-category Switch trials (24). The order of the trials was determined pseudo-randomly while balancing the types of switches. Each run included a total of 73 trials, with an extra trial at the beginning of the run that was required for the balancing of the switch types. This trial was assigned a random rule and was excluded from the analysis.

To address multivariate effects, we used decoding of rule pairs in the task. Each pair of rules out of the six could then be referred to as either ‘within-category’, i.e., applied on the same category domain, or ‘between-category’, i.e., applied on different category domains. The idea being, while all rules may be decoded across the frontoparietal network, the ‘between-category’ rules might be more distinct (i.e. higher decoding levels) than ‘within-category’ rules (Crittenden et al., 2016). To avoid confounding the rule representation with visual information in the task as in Crittenden et al. 2016, we used a variant of this task, with separate cue and stimulus presentation phases in each trial (Smith et al., 2018).

The participants practiced the task prior to the scanning session until they learned the rules. The practice consisted of two parts. During the first part, trials included feedback, while feedback was omitted in the second part. There was no feedback during the scanning session runs of this task. During the scanning session, after completing the localizer tasks, the participants were asked to state the six rules to make sure they remembered the rules before starting the rule-based task. They were also shown the rules again if they requested it. The task was self-paced, with an average duration of each run of 5 minutes and 52 seconds (SD = 29.1 seconds).

### 2.4. fMRI data acquisition

Participants were scanned in a Siemens 3T Prisma MRI scanner with a 32-channel head coil. A T2*-weighted 2D multiband Echo-Planar Imaging, with a multi-band factor of 3, was used to acquire 2 mm isotropic voxels (Feinberg et al., 2010). Other acquisition parameters were: 48 slices, no slice gap, TR = 1.1 s, TE = 30 ms, flip angle = 62⁰, field of view (FOV) = 205 mm, in plane resolution: 2 x 2 mm. In addition to functional images, T1-weighted 3D multi-echo MPRAGE (van der Kouwe et al., 2008) structural images were obtained (voxel size 1 mm isotropic, TR = 2530 ms, TE = 1.64, 3.5, 5.36, and 7.22 ms, FOV = 256 mm x 256 mm x 192 mm). A single structural image was computed per subject by taking the voxelwise root mean square across the four MPRAGE images that are generated in this sequence.

### 2.5. Data Analysis

#### 2.5.1. Pre-processing

Pre-processing was performed using the automatic analysis (aa) pipelines (Cusack et al., 2014) and SPM12 (Wellcome Trust Centre for Neuroimaging, University College London, London) for MatLab. The data were first motion corrected by spatially realigning the EPI images. The images were then unwarped using the fieldmaps, slice time corrected and co-registered to the structural T1-weighted image. The structural data were normalized to the Montreal Neurological Institute (MNI) template using nonlinear deformation, after which the transformation matrix was used to normalize the EPI images. For the univariate ROI analyses, the localizer data were pre-processed without smoothing. For the whole-brain random-effects analysis and voxel selection for subject-specific ROIs, the localizer data were smoothed with a Gaussian kernel of 5 mm full-width at half-maximum (FWHM). No smoothing was applied to the data of the rule-based criterion task, for both the multivariate and univariate ROI analyses, thus assuring that identical data is used for both analyses.

#### 2.5.2. General linear model (GLM)

A general linear model (GLM) was estimated per participant for each localizer task. Regressors were created for the Easy and Hard task blocks and were convolved with a canonical hemodynamic response function (HRF). Run means and movement parameters were used as covariates of no interest. The resulting β-estimates were used to construct the contrast of interest between the Hard and Easy conditions, and the difference in β-estimates (Δbeta) was used to estimate the activation evoked by each localizer.

For the rule-based criterion task, two different GLMs were used to address univariate and multivariate effects. The GLM for the univariate analysis was based on types of switches between trials. A GLM was estimated for each participant using cue phase regressors for the three switch types: Stay, within-category Switches and between-category Switches. Additional stimulus phase regressors were used separately for each switch type, from stimulus onset to response, but these were not analyzed further. A similar GLM was used for multivariate analysis, based on the six rule types instead of switch types. The duration of the cue regressors was 2 s for all conditions in both models. The regressors were convolved with the canonical HRF and the six movement parameters and run means were included as covariates of no interest.

#### 2.5.3. Localizers activity patterns

We used both random-effects whole-brain and ROI analyses to assess and compare the activation levels and recruitment of the MD network by the localizers. All measures used voxel data of the Hard versus Easy contrast and the resulting difference in beta estimates (Δbeta), computed across the two runs of each localizer as well as separately for each run when required as detailed below for the specific analyses. Additional Hard versus Easy contrasts were computed using combinations of different localizers.

For ROI analysis, we used a template for the MD network, derived from an independent task-based fMRI dataset (Fedorenko et al., 2013, http://imaging.mrc-cbu.cam.ac.uk/imaging/MDsystem). The template is bilateral with an equal number of voxels in each hemisphere for each ROI. ROIs and their respective number of voxels per hemisphere include the anterior insula (AI, 992 voxels), posterior/dorsal lateral frontal cortex (pdLFC, 1132 voxels), intraparietal sulcus (IPS, 4260 voxels), the anterior, middle and posterior middle frontal gyrus (MFG) (621, 712 and 1269 voxels, respectively) and the pre-supplementary motor area (preSMA, 1247 voxels). The group-level ROI analyses were computed using the MarsBaR SPM toolbox (Brett, Anton, Valabregue, & Poline, 2002).

#### 2.5.4. Measures to compare activation patterns of the localizers

We conducted several analyses to compare the spread and similarity of activation patterns of the localizers.

##### 2.5.4.1 Whole-brain spatial spread of activity patterns of localizers across subjects

We conducted a whole-brain analysis to examine the spread of activation patterns across individual participant data. For each voxel we computed the number of participants with significant activations by applying FDR (p < 0.05) across all voxels and all participants. This yielded a whole-brain map in which voxel data represents the number of participants with significant activation.

##### 2.5.4.2 Correlation measures to compare activation patterns of the localizers

To quantify and compare activity patterns of the three localizers, we used Fisher-transformed Pearson correlations of Δbeta estimates (Hard – Easy) across voxels. Contrast data were estimated for each run separately and correlations were computed across all voxels in each MD ROI. First, we computed the reliability of activation patterns. For each subject and localizer we correlated the Δbeta estimates of the two runs, separately for MD ROIs and then averaged. Second, to compare activity patterns between subjects, we then estimated the similarity in activity patterns between subjects. For each subject, each localizer, and each MD ROI, we computed the correlation of Δbeta estimates between the first run of that subject and the second run of another subject of the same localizer. For each subject, this was computed 17 times using all other subjects and averaged across them to get between-subject correlations of activity, separately for each of the three localizers. Lastly, to estimate the similarity in activity patterns between the different localizers, we computed for each subject the correlation between the first runs of each pair of localizers. Only one run of each localizer was used to estimate these between-localizer similarities in order to keep the analysis and resulting correlations as similar as possible to the within-subject within-localizer reliabilities and to allow the comparison between the two. While Fisher-transformed correlations were used for statistical inference, Pearson correlations are presented in the text and figures for ease of interpretation.

#### 2.5.5. Individual MD localization using the group-constrained subject-specific approach

Individual subject ROIs were defined using the group-constrained subject-specific approach. For each localizer, we used the Hard versus Easy contrast data across the two runs to obtain Δbeta estimates. Then, for each ROI, the 200 voxels with the largest Δbeta estimates were selected. The number of voxels that were selected was defined prior to any data analysis.

The selected voxels were then used to test for the effects of subject-specific ROIs and choice of localizer on both the univariate and multivariate activity as measured in the rule-based criterion task. We compared measures of activity using both subject-specific ROIs based on the different localizers and the group template (i.e., using all voxels within each ROI), as well as between localizers.

#### 2.5.6 Univariate analysis of the criterion task

For the univariate activity in the criterion task, we used two contrasts with varying cognitive demand to test for the effect of subject-specific ROIs and choice of localizer across the MD network. For each subject, the GLM cue regressors across all four runs were used to compute two univariate contrasts: within-category Switch versus Stay trials, and between-category Switch versus Stay trials. The results were then averaged across hemispheres and subjects. Similarly to the Hard vs. Easy contrast in the localizer tasks, activity across the MD network is expected to increase with increased demand.

#### 2.5.7. Multivoxel pattern analysis (MVPA) of the criterion task

We used rule decoding in the criterion task to test for the effect of subject-specific ROIs and choice of localizer on multivariate activity. The decoding analysis focused on the task rules during the cue phase, when only colored frames appear on the screen, therefore avoiding any confounds of the subsequent stimuli. Classification accuracy was computed using a support vector machine classifier (LIBSVM library for MATLAB, c=1) implemented in the Decoding Toolbox (Hebart, Gorgen, & Haynes, 2015). A leave-one-run-out cross-validation was employed to compute pairwise classifications for all task rule combinations (15 in total), and classification accuracy was averaged across all folds for each pair of rules. The average accuracy of all rule pairs, as well as separately for within- and between-category rule pairs, was computed for each subject and ROI. For each subject, classification accuracies were computed using subject-specific ROIs based on the three independent localizers’ data separately, as well as using all voxels within each ROI (i.e. group template). Decoding accuracies above chance (50%) were then averaged across hemispheres and subjects and were tested against zero using one-tailed t-tests. As argued by (Allefeld, Görgen, & Haynes, 2016), such a t-test implements a fixed effects analysis, which is suitable for our systematic comparison of decoding results in a given criterion task when using different localizers or all voxels to define ROIs.

Since all localizers were chosen because their robust and consistent recruitment of the MD network, combinations of localizers could also be used to define subject-specific ROIs. To test for multivariate results in subject-specific ROIs defined using data combined from two different localizers, we repeated the decoding analysis using contrast data comprised of two runs, one of each localizer. We used the first run of each localizer and created three combination contrasts from pairs of localizer tasks: spatial WM + verbal WM, spatial WM + Stroop, and verbal WM + Stroop. We also used all 6 localizer runs to create a spatial WM + verbal WM + Stroop contrast. The combination contrasts were used to define subject-specific ROIs for the decoding analysis in a similar way to the individual localizers’ data.

One advantage of using group-constrained subject-specific ROIs for multivariate analysis in particular is that it allows keeping the number of voxels in each ROI fixed and controlled. This may be important as it has been previously demonstrated in the visual system that the size of ROI may affect decoding accuracy levels (Eger et al., 2008; Said et al., 2010; Walther et al., 2009). To test for potential effect of ROI size on the decoding results, the MVPA was repeated using a range of ROI sizes (50, 100, 150, 200, 250, 300, 350 and 400 voxels).

To ensure that our decoding results did not depend on the choice of classifier, we repeated the MVPA using a representational similarity analysis (RSA) approach, with linear discriminant contrast (LDC) as a measure of dissimilarity between rule patterns (Carlin & Kriegeskorte, 2017; Nili et al., 2014). Cross-validated Mahalanobis distances were calculated for all 15 pairwise rule combinations and averaged to get within- and between-category rule pairs, for each ROI and participant, using all the voxels in the ROI and subject-specific ROIs defined using the different localizer tasks. For each pair of conditions, we used one run as the training set and another run as the testing set. This was done for all pairwise combinations of the 4 runs and LDC values were then averaged across them. Larger LDC values indicate more distinct patterns of the tested conditions, while the LDC value itself is non-indicative for level of discrimination. The choice of using LDC rather than LD-t (associated t-value) meant that we could meaningfully look at differences between distances, and particularly the distinction between within- and between-category rule discriminations in the criterion task. We therefore used the difference of between- and within-category rule pairs compared to 0 as indication for representation of rule information.

#### 2.5.8 Statistical testing and code

To compare performance of the localizers to the group template as well as to each other, we first used a repeated measures ANOVA as a statistical model with factors as appropriate for each question and as detailed in the Results. The main factor of interest in all these ANOVAs is the localizer factor (with levels for the three localizers and the group template when required), and we report all the results related to this factor and its interactions, but not other interactions that are not of interest for our questions. To directly test for our research questions, we also used separate t-tests for each localizer compared to the group template when appropriate, as well as for all possible pairs of localizers, both corrected for multiple comparisons. For both univariate and multivariate data, the main question of interest when comparing the localizers and the group template is whether using individual ROIs affects the power to detect an underlying effect. Increased power can be achieved either by increases in the means or by reduced between-subject variability. Therefore, in addition to tests for mean differences using an ANOVA as detailed above, we also tested for differences in variance in our main analyses. We used Mauchly’s test of sphericity (using SPSS V25) applied on the appropriate repeated-measures model followed by post-hoc pairwise comparisons using Pitman-Morgan test for equality of variance in dependent samples when required.

We used an alpha level of .05 for all statistical tests. Bonferroni correction for multiple comparisons was used when required, and the corrected p-values and uncorrected t-values are reported. To quantify the evidence for differences in pattern discriminability when using the group template and subject-specific ROIs defined by functional localizers, we conducted a complementary Bayes factor analysis (Rouder et al. 2009). We used JZS Bayes factor for one-sample *t*-test and square-root(2)/2 as the Cauchy scale parameter, therefore using medium scaling. The Bayes factor is used to quantify the odds of the alternative hypothesis being more likely than the null hypothesis, thus enabling the interpretation of null results. A Bayes factor greater than 3 is considered as some evidence in favor of the alternative hypothesis. All analyses were conducted using custom-made MATLAB (The Mathworks, Inc) scripts, unless otherwise stated.

### 2.6. Data and code availability statement

Anonymized data and code will be available in a public repository before publication. Data and code sharing are in accordance with institutional procedures and ethics approval.

## 3. Results

### 3.1. Behavioral results

The mean accuracies and reaction times (RT) for the Easy and Hard conditions of the spatial WM, verbal WM, and Stroop localizer tasks are listed in Table 1. As expected, there was a significant increase in RT during the Hard compared to the Easy condition for all localizers (two-tailed paired t-test: spatial WM: *t_17_* = 10.03, *p* < 0.001; verbal WM: *t_17_* = 25.01, *p* < 0.001; Stroop: *t_17_* = 9.89, *p* < 0.001), as well as a significant decrease in accuracy (spatial WM: *t_17_* = 8.65, *p* < 0.001; verbal WM: *t_17_* = 6.95, *p* < 0.001; Stroop: *t_17_* = 7.47, *p* < 0.001). These results confirmed that the task manipulation of Easy and Hard conditions worked as intended.

**Table 1.** RTs and accuracies in the localizer tasks. Values are means ± standard errors.

|  | Spatial WM |  | Verbal WM |  | Stroop |  |
| --- | --- | --- | --- | --- | --- | --- |
|  | Easy | Hard | Easy | Hard | Easy | Hard |
| RT (sec) | 1.15±0.05 | 1.63±0.06 | 1.02±0.04 | 1.91±0.04 | 0.78±0.02 | 0.90±0.02 |
| Accuracy<br>(% correct) | 94.2±1.3 | 71.4±2.5 | 96.8±0.9 | 79.2±2.6 | 94.4±0.9 | 68.5±4.1 |

Accuracy levels for the rule-based criterion task were high (mean ± SD: 95.3% ± 2.3), indicating that the participants were able to learn the different rules and apply them correctly. The mean accuracies (mean ± SD): 93.5±3.4, 95.7±3.2, 94.1±4.2, 95.4±3.3, 95.4±4.6, 97.7±3.0, and the mean RTs (mean ± SD): 1.33±0.4 s, 1.46±0.5 s, 1.55±0.5 s, 1.55±0.6 s, 1.27±0.4 s, 1.30±0.3 s, for age, gender, building type, viewpoint, first letter, and last letter rules, respectively. Since the focus of this study was to compare imaging results across different methods rather than linking them to behavior or making inference about the underlying cognitive construct, we did not analyze the accuracy levels and the RTs for the criterion task any further.

### 3.2 Whole-brain and ROI univariate analysis of the localizer tasks

To test for the recruitment of the MD network, a whole-brain random effects analysis was conducted for the Hard versus Easy contrasts of each localizer task (**Figure 2A**). The whole-brain patterns of activity clearly showed recruitment of the MD network by all localizers. Areas of increased activity included the anterior-posterior axis along the middle frontal gyrus, anterior insula, and the area anterior to the FEF on the lateral surface; preSMA on the medial surface; and IPS on both the lateral and medial surfaces. An additional visual component was observed, as expected from the nature of the tasks. The pattern of activity for the Stroop localizer was sparser, in particular on the right hemisphere.

**Figure 2.**
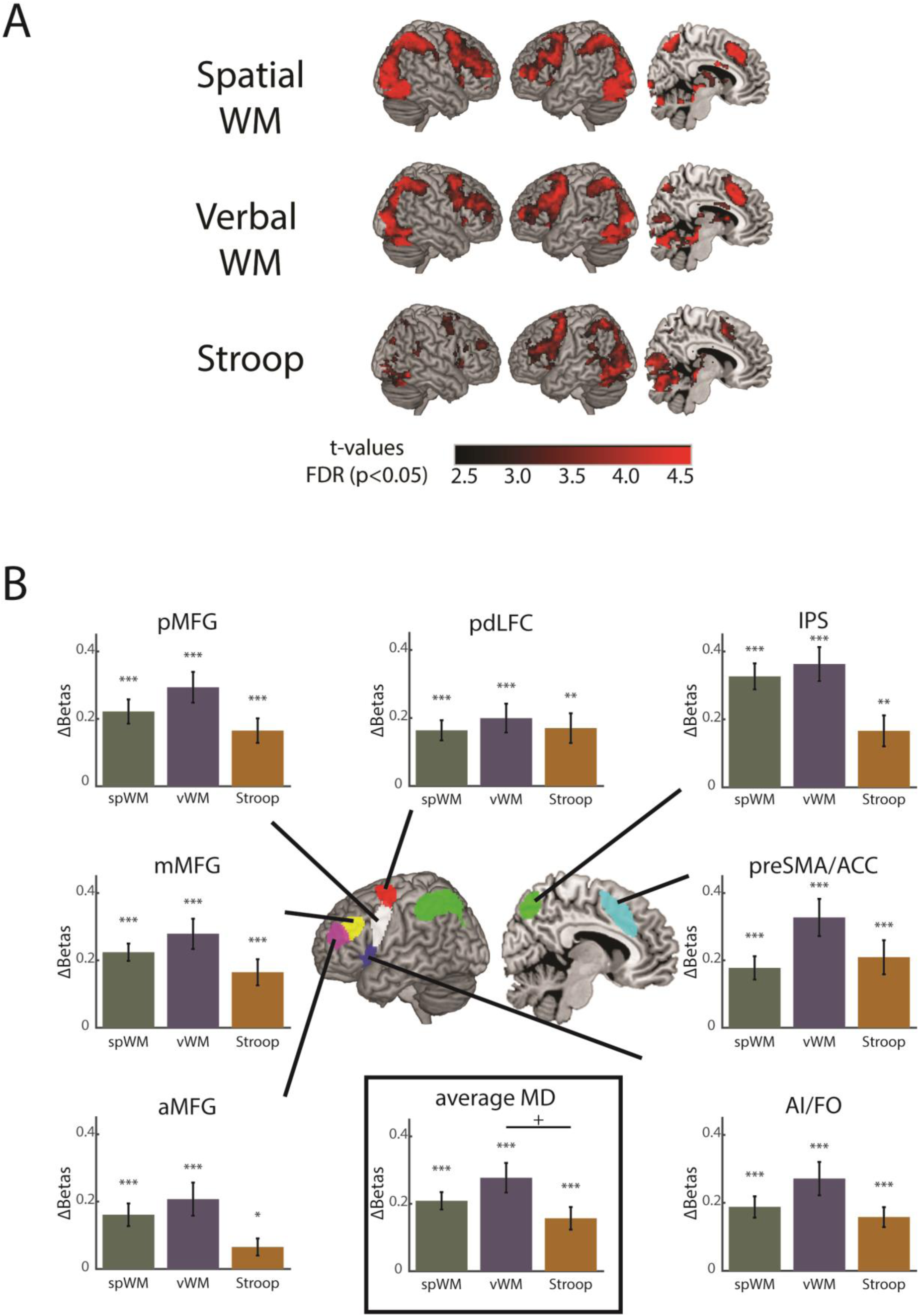
Increased activity across the MD network with increased difficulty level for all three localizers. **A:** Whole-brain t-maps of the contrast between Hard and Easy conditions for the localizer tasks. WM = working memory. t-maps are FDR corrected, p<0.05. **B:** Univariate results for the contrast between the Hard and Easy conditions for the three localizer tasks, per MD ROI and averaged across ROIs. Significance levels above zero in each ROI and for each localizer (with Bonferroni correction for 3 comparisons) are shown to demonstrate recruitment of the MD network. For the average across MD ROIs, pairwise statistical testing between localizers was also done using a paired two-tailed t-test with Bonferroni correction (3 comparisons). Error bars indicate SEM. The MD template that was used for ROI analysis is shown in the middle for reference (adapted from Fedorenko et al. 2013). spWM = spatial working memory, vWM = verbal working memory. pdLFC = posterior/dorsal lateral prefrontal cortex, IPS = intraparietal sulcus, preSMA = pre-supplementary motor area, AI = anterior insula, aMFG = anterior middle frontal gyrus, mMFG = middle frontal gyrus, pMFG = posterior middle frontal gyrus. * *p* < 0.05, ** *p* < 0.01, *** *p* < 0.001, + *p* < 0.06.

An ROI analysis further confirmed the recruitment of the MD network with increased task difficulty (**Figure 2B**). All localizers showed a significant increase in activation for the Hard compared to the Easy condition in each of the ROIs with Bonferroni correction (3) for multiple comparisons (one-sample two-tailed t-test against 0: *t_17_* ≥ 2.57, *p* ≤ 0.02 for all). A three-way repeated measures ANOVA with factors task (spatial WM, verbal WM, Stroop), ROI (7) and hemisphere (left, right) revealed a main effect of task (*F_2, 34_* = 3.96, *p* = 0.028). There was a significant main effect of ROIs (*F_6, 102_* = 7.6, *p* < 0.001), indicating some differences in activity levels between the ROIs. Although the recruitment of the two hemispheres was broadly similar, there was also an effect of hemisphere (*F_1, 17_* = 15.54, *p* = 0.001), with larger activity on the left than the right hemisphere. There were significant interactions between tasks and ROIs, tasks and hemispheres, ROIs and hemispheres and a three-way interaction (*F* > 4.1 *p* < 0.009). Post-hoc pairwise comparisons with Bonferroni correction (3 comparisons) across ROIs demonstrated a marginal difference in activity between the verbal WM and Stroop task (*t_17_* = 2.64, *p* = 0.051), but the overall activity did not differ between the spatial WM and verbal WM tasks or the spatial WM and Stroop tasks (*t_17_* ≤ 1.25, *p* ≥ 0.3 for both). Overall, all the tasks recruited the MD network, with some differences in the activation patterns across ROIs for the different localizers. Importantly, the univariate results confirmed a significant increase in activity in the MD network with increased task difficulty for all three localizers, as expected and designed.

### 3.3 Comparisons of activity patterns between localizers

We further quantified and compared the variability of activity patterns across subjects and localizers. To test for variability across subjects, two methods were used: a whole-brain overlay of subjects’ activation per voxel and correlations of activity across voxels between runs of the same subject versus different subjects. Variability across localizers was tested by correlating the activity across voxels between runs of the same localizer versus different localizers.

#### 3.3.1 Variability of activity patterns across subjects

While the group averages of the Hard versus Easy contrasts of the three localizers closely resembled the MD template, there was substantial variability between activation patterns of individual participants. To visualize this spread of activations across individuals, we computed a whole-brain overlay map for each localizer (**Figure 3A**). Each voxel shows the number of participants with significant activations after applying FDR (p < 0.05) correction across voxels and subjects. The overlay maps show peaks of activation for many subjects in areas similar to those observed in the group averages. However, it is also clear from these maps that there is substantial variability across subjects with activations that extend to large parts of the frontal and parietal cortices, as well as the visual cortex as expected from the visual nature of the tasks.

**Figure 3.**
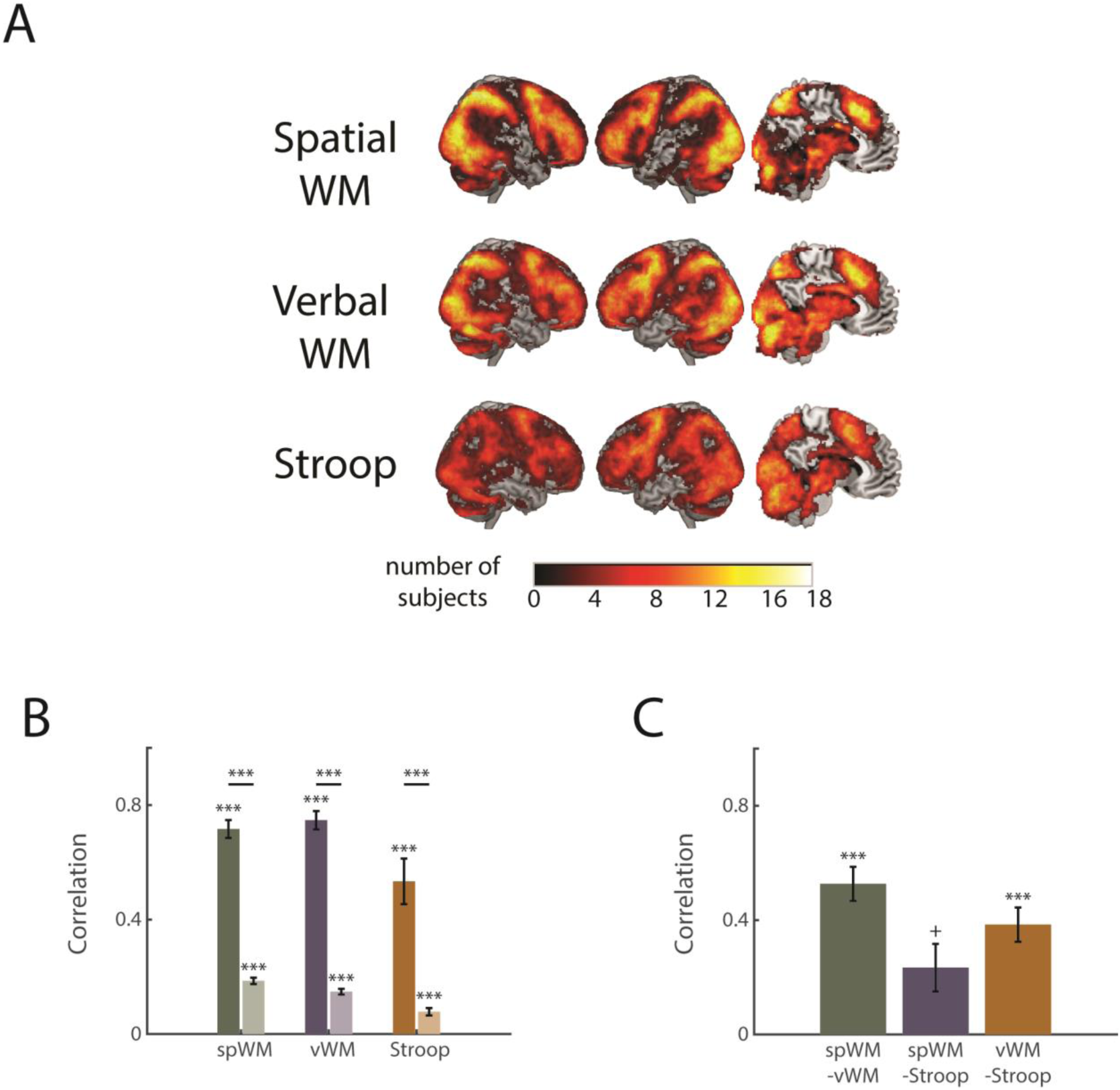
Activity patterns are variable across subjects and localizers. **A:** Activation patterns of single subjects only partially overlap, demonstrating variability across participants. The color of each voxel shows the number of subjects that had significant activation in that voxel in the Hard versus Easy contrast, with the color bar indicating number of subjects up to 18 (sample size), thresholded at 1 subject. **B.** Correlation of activity within runs of the same subject (reliabilities, darker bars) and runs of different subjects (lighter bars) for each localizer, averaged across MD ROIs. **C.** Within-subject correlation of activity patterns between pairs of localizers, averaged across MD ROIs. Pearson correlations are presented, while Fisher transformed correlations were used for statistical inference. Asterisks above bars show significance levels (two-tailed t-test against zero, corrected for 3 comparisons). Asterisks above horizontal lines between bars show significance levels of differences (paired two-tailed t-test against zero, corrected for 3 comparisons). Error bars indicate SEM. * *p* < 0.05, ** *p* < 0.01, *** *p* < 0.001, + *p* < 0.06.

Correlations of activity of the different localizers for each subject and between subjects further demonstrated that activity patterns are subject-specific. For each localizer, we computed the reliability of activation as the correlation between runs of the same localizer and same subject using all voxels within each ROI of the MD template. These reliabilities were compared to correlation between two runs from different subjects and same localizer. The correlations were averaged across ROIs to get the correlations of activity across the MD network (**Figure 3B**). The reliability of activation patterns for each subject was high for all localizers (Mean ± SD: spatial WM: 0.72 ± 0.13; verbal WM: 0.75 ± 0.14; Stroop: 0.53 ± 0.34), and their similarity was substantially lower for different subjects (Mean ± SD: spatial WM: 0.19 ± 0.05; verbal WM: 0.15 ± 0.04; Stroop: 0.08 ± 0.06). A two-way repeated measures ANOVA with task (3: spatial WM, verbal WM and Stroop) and correlation type (2: within- or between-subjects) as factors showed that the within-subject reliabilities were significantly larger than the between-subject correlations (*F_1, 17_* = 265.9, *p* < 0.001), providing additional support for the large variability of activity patterns across subjects. There was also a significant main effect of task (*F_2, 34_* = 7.5, *p* < 0.01), and post-hoc pairwise comparisons with Bonferroni correction (3 comparisons) showed that the Stroop task had lower correlations than the verbal WM task (*t_17_* = 3.42, *p* = 0.01) and a trend towards lower correlation compared to the spatial WM task (*t_17_* = 2.6, *p* = 0.062). There was also a marginal interaction between task and correlation type (*F_2, 34_* = 3.2, *p* = 0.053), but within-subject reliabilities were larger than between-subject correlations for all three localizers (*t_17_* > 6, *p* = 0.001).

#### 3.3.2 Variability of activity patterns across localizers

To assess the similarity of activity patterns between localizers, we computed the within-subject correlation of activity pattern across voxels between the first runs of pairs of localizers (**Figure 3C**). As expected, there was a substantial positive correlation between the localizers (Mean ± SD: spatial WM with verbal WM: 0.53 ± 0.25; spatial WM with Stroop: 0.23 ± 0.35; verbal WM with Stroop: 0.38 ± 0.25), with correlations between the working memory tasks and between verbal WM and Stroop being well above 0 (two-tailed t-test against zero corrected for 3 comparisons: *t* > 6, *p* < 0.001), and a marginally significant correlation between the spatial WM and Stroop correlation following Bonferroni correction (*t*_17_ = 2.65, *p* = 0.051). These correlations were significantly smaller than the within-subject within-localizer reliabilities (two-tailed t-test of the average reliabilities across localizers vs. the average correlations of pairs of localizers: *t_17_* = 9.37, *p* < 0.001). These reduced correlations demonstrated that although all localizers recruited the MD network, there was some spatial variability in their activation patterns.

### 3.4 Subject-specific ROIs and univariate activity in the rule-based criterion task

The main aim of our study was to test for the effect of subject-specific ROIs on univariate and particularly multivariate activity measures in the MD network. We used the rule-based criterion task to extract both univariate and multivariate measures and tested whether using subject-specific ROIs using the independent localizers’ data affects the ability to identify activity as expected in the MD network, and whether such changes depend on the choice of localizer.

For the univariate activity, we computed two task switch contrasts in the criterion task: the more demanding between-category Switch versus Stay trials, and the less demanding within-category Switch versus Stay trials. The analysis was done using subject-specific ROIs based on the localizers’ data as well as using all voxels within each ROI and averaged across all MD ROIs (**Figure 4**). A three-way repeated measures ANOVA with task (spatial WM, verbal WM, Stroop, all-voxels), contrast type (within-category Switch versus Stay, between-category Switch versus Stay), and ROI (7) was set to test for the effect of using subject-specific ROIs and localizer choice on mean activity levels. There was a main effect of task (*F_3, 51_* = 14.21, *p* < 0.001), and post-hoc tests with Bonferroni correction for 6 comparisons showed that activity when using the group template was lower than when using subject-specific ROIs, for all localizers (*t_17_* > 3.5, *p* < 0.02). The activation using subject-specific ROIs based on the spatial WM localizer was lower than that of verbal WM (*t_17_* = 3.40, *p* = 0.02) and other comparisons between localizers were not significant (*t_17_* > 1.4, *p* > 0.9). As expected for activity in the MD network, there was a main effect of contrast type (*F_1, 17_* = 12.12, *p* = 0.003), with activity for the more demanding between-category Switch being larger than the within-category Switch. There was also an interaction between task and contrast type (*F_3, 51_* = 7.92, *p* = 0.002), but post-hoc tests with Bonferroni correction for 4 comparisons showed that activity for the between-category Switch was larger than for the within-category Switch for all localizers and when the group template was used (*t_17_* > 2.9, *p* < 0.036). Overall, these results demonstrate that using subject-specific ROIs leads to an increase in the observed mean univariate results, with similar effects for the different localizers.

**Figure 4.**
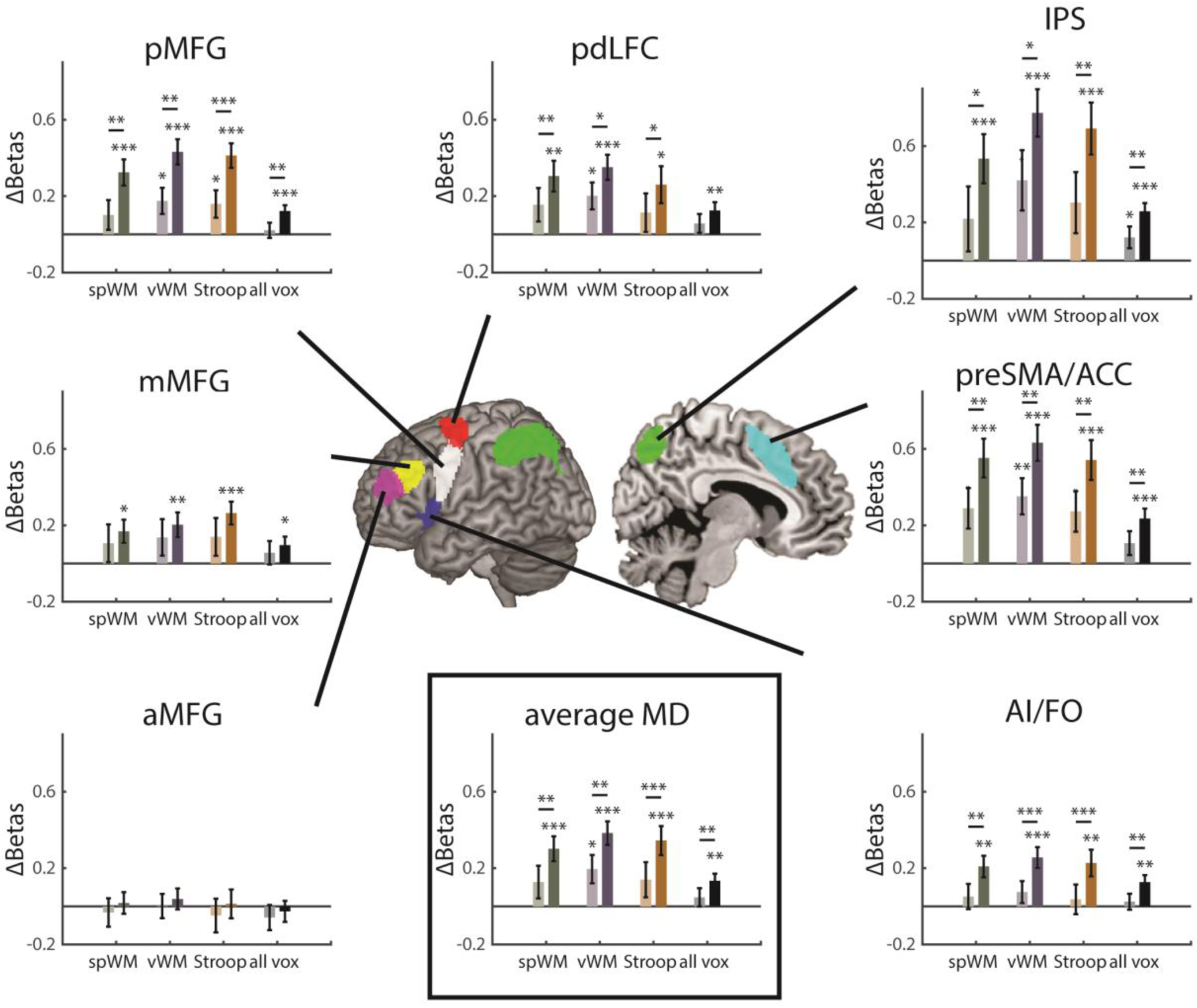
Univariate activity in the criterion task increases when using subject-specific ROIs compared to the group template. Within-(lighter bars) and between-(darker bars) category versus Stay trials contrast values using subject-specific ROI defined by the different localizer tasks and using all voxels (group template, no localizer), per ROI and averaged across MD ROIs. Δbeta values were larger when using subject-specific ROIs compared to when using the group template, and similar for all three localizers. Between-category Switch activity was larger than the within-category Switch for all localizers and when using all voxels, as expected for the MD network. spWM = spatial working memory, vWM = verbal working memory. pdLFC = posterior/dorsal lateral prefrontal cortex, IPS = intraparietal sulcus, preSMA = pre-supplementary motor area, AI = anterior insula, aMFG = anterior middle frontal gyrus, mMFG = middle frontal gyrus, pMFG = posterior middle frontal gyrus. Significant activation levels are shown above bars (two-tailed t-test against 0). For visualization, asterisks above horizontal lines between bars show significance levels of differences (paired two-tailed t-test against zero). Error bars indicate SEM. * p < 0.05, ** p < 0.01, *** p < 0.001.

In addition to the mean differences that we observed, differences in between-subject variability may contribute to the power to detect an underlying effect. To test for differences in variability when individual ROIs are used and for the effect of choice of localizer, we applied a Mauchly’s test of sphericity on the same repeated measures model that was used for the test of means, with factors task (spatial WM, verbal WM, Stroop, all-voxels), contrast type (within-category Switch versus Stay, between-category Switch versus Stay), and ROI (7). Of particular interest to our question were effects of localizer and an interaction between localizer and contrast type, and both showed a significant difference of variance (*W_5_* = 0.42, *p* = 0.018; *W_5_* = 0.27, *p* = 0.001, respectively). Post-hoc Pitman-Morgan tests of all pairwise tasks (corrected for 6 comparisons) on the difference in ΔBeta between between-category and within-category Switches showed no differences in variance between the three localizers (*t_17_* < 1.9, *p* > 0.4). However, the variance of the univariate activity was larger when using all voxels in the group template compared to using individual ROIs, for all three localizers (*t17* > 5.18, *p* < 0.001), possibly simply because univariate activity is an average across more voxels when all voxels are used.

### 3.5 Subject-specific ROIs and rule decoding in the rule-based criterion task

We next tested for the effect of subject-specific ROIs on decoding levels in the rule-based criterion task across the MD network. We computed the decoding accuracy of all pairwise discriminations between rules when using each localizers’ data to select the 200 most responsive voxels within each ROI for each subject and when using all the voxels in each ROI. Overall decoding accuracies above chance (50%) across all MD ROIs and rule-pairs were 2.85 ± 14.53, 2.71 ± 14.75, 3.17 ± 15.04, 5.08 ± 15.34 for subject-specific ROIs defined using the spatial WM, verbal WM and Stroop localizers, and when using the group template, respectively (mean ± SD). These decoding levels were similar to previous studies that used a similar experimental paradigm (Crittenden et al., 2015; Smith et al., 2018). We first tested whether overall mean decoding accuracy across all pairs of rules differed between the localizers and compared to when all voxels within each ROI were used. A two-way repeated measures ANOVA with task (spatial WM, verbal WM, Stroop, all-voxels) and ROI (7) showed a main effect of task (*F_3, 51_* = 2.9, *p* = 0.042). However, none of the post-hoc tests to compare pairs of tasks survived correction (Bonferroni correction for 6 comparisons, *t_17_* < 2.7, *p* > 0.09). There was a main effect of ROI (*F_6, 102_* = 2.5, *p* = 0.025) but no interaction with task (*F_18, 306_* = 0.8, *p* = 0.7). Overall, decoding across all pairs of conditions was similar for all localizers and when the group template was used.

Our choice of criterion task has enabled us to not only test for the effect of localizer type on the overall discrimination between rules, which might be too coarse to depict, but also for the relative decoding levels of the two types of discriminations, thus potentially picking up more subtle effects. The criterion task included discriminations between rules applied on the same category (within-category discriminations, e.g., between the gender and age rules, both applied on the faces category) and discriminations between rules applied on different categories (between-category discriminations, e.g., between the gender rule applied on the faces category and the viewpoint rule applied on the building category). To get this more fine-grained picture of the effect of subject-specific ROIs using localizer data on rule decoding, as well as when the groups template is used, we split the decoding accuracy to within- and between-category rule discriminations (**Figure 5**). A three-way repeated measures ANOVA with task (spatial WM, verbal WM, Stroop, all-voxels), distinction type (within-category, between-category) and ROI (7) as within-subject factors was set to test for differences between localizers related to mean decoding of the two distinction types. There was no main effect of task (*F_3, 51_* = 1.0, *p* = 0.4), and no interaction between distinction type and task (*F_3, 51_* = 0.9, *p* = 0.4). To test for differences between each of the localizers and the group template as well as pairs of localizers based on our pre-defined questions, additional t-tests between all pairs of tasks were conducted, but no significant differences were found (*t_17_* < 1.8, *p* > 0.5, corrected for 6 comparisons). There was no main effect of distinction type (*F_1, 17_* = 3.2, *p* = 0.089), though a numerical trend was consistent with the previously reported results (Crittenden et al., 2016). There was no main effect of ROI or interaction of ROI with distinction type or task (*F* < 1.8, *p* = 0.09). Taken together, this indicates that decoding results in the criterion task were similar for all localizers as well as when the group template was used, similar to the results obtained across all pairs of conditions.

**Figure 5.**
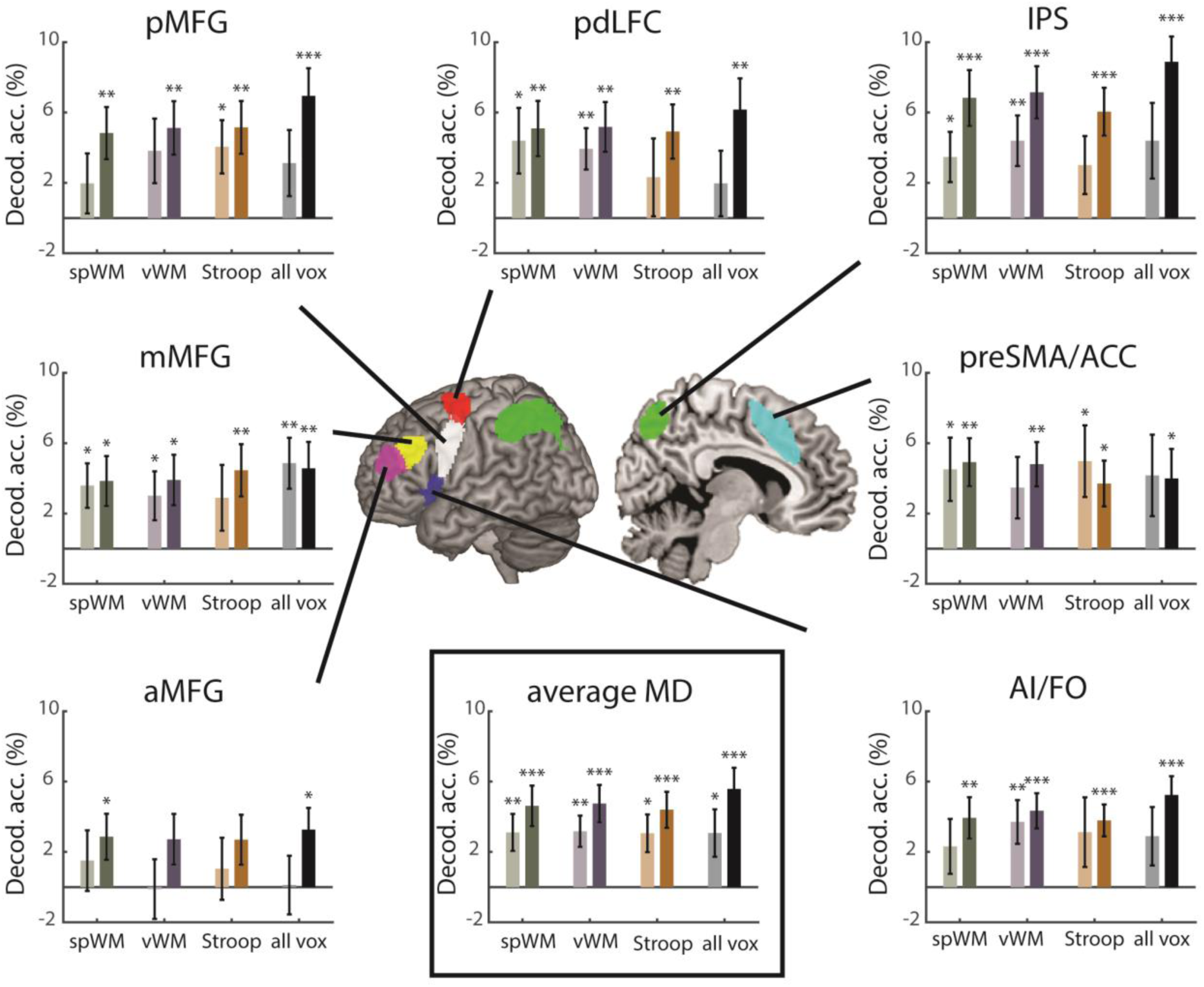
Decoding accuracy for within- and between-category rule pairs. Within-(lighter bars) and between-(darker bars) category rule decoding accuracy values above chance (50%) for subject-specific ROIs defined using the different localizer tasks and using all voxels (group template), per ROI and averaged across MD ROIs. Decoding accuracies were similar for all three localizers and when using all voxels within each template ROI, with similar differences between within- and between-category rule decoding (see Text (*3.5*) for statistical details). spWM = spatial working memory, vWM = verbal working memory. pdLFC = posterior/dorsal lateral prefrontal cortex, IPS = intraparietal sulcus, preSMA = pre-supplementary motor area, AI = anterior insula, aMFG = anterior middle frontal gyrus, mMFG = middle frontal gyrus, pMFG = posterior middle frontal gyrus. To demonstrate rule decoding, significant decoding accuracy above chance (50%) is shown (one-tailed t-test against 0). Error bars indicate SEM. * p < 0.05, ** p < 0.01, *** p < 0.001.

Based on these ANOVA results and to further establish that using localizer data for subject-specific ROIs did not change decoding levels, we conducted a Bayes factor analysis (Rouder et al. 2009), separately for each functional localizer compared to decoding with all voxels. First, the difference in classification accuracy between the between- and within-category distinctions averaged across ROIs for each functional localizer was compared to the classification accuracies with all voxels using a paired two-tailed t-test. The t-value was then entered into a one-sample Bayes factor analysis with a Cauchy scale parameter of 0.7. The Bayes factors for the spatial WM, verbal WM and Stroop localizer when compared to the decoding levels with all voxels were 0.40, 0.56 and 0.79, respectively. These results demonstrate little evidence for differences in decoding levels when subject-specific ROIs and the group template are used.

Beyond changes in mean decoding levels, we further tested for differences in between-subject variability that may affect the power to detect an underlying change in decodability when subject-specific ROIs are used. A Mauchly’s test for sphericity applied on the same repeated measures model that was used to test for mean differences with factors task (spatial WM, verbal WM, Stroop, all-voxels), distinction type (within-category, between-category), and ROI (7) showed no effect of localizer (*W_5_* = 0.54, *p* = 0.09) and no interaction between localizer and distinction type (*W_5_* = 0.67, *p* = 0.27). Overall, we concluded that using subject-specific ROIs did not change neither the mean decoding levels nor the between-subject variability, with similar results for all three localizers.

Individual-subject localization of the MD network for MVPA can be done not only using one localizer task, but also using two runs of two different localizers, thus benefitting from the activation patterns evoked by both localizers to more robustly identify MD-like voxels. To test for the effect of such an approach on decoding results, we repeated the analysis using subject-specific ROIs based on localizer data combined from two different localizers (spatial WM + verbal WM, spatial WM + Stroop, verbal WM + Stroop). A three-way repeated measures ANOVA with task (4: three combination contrasts and all-voxels), distinction type (within/between category) and ROI (7) as within-subject factors did not show a main effect of task (*F_3,_* _51_ = 2.5, *p* = 0.07), similarly to the results when individual localizers were used to define subject-specific ROI. There was no effect of distinction type (*F_1, 17_* = 3.3, *p* = 0.08), despite a numerical trend, and there was no interaction of task and distinction type (*F_3, 51_* = 1.4, *p* = 0.25). Similarly, subject-specific ROIs were also defined using all localizers with one contrast using all 6 localizer runs (spatial WM + verbal WM + Stroop). A three-way repeated measures ANOVA with task (2: one combination contrast and all-voxels), distinction type (within/between category) and ROI (7) as within-subject factors showed no main effect of task or distinction type or an interaction of the two (*F* < 3.2, *p* > 0.05). Overall, these results indicate that using combinations of localizer tasks to define subject-specific ROIs yielded decoding results similar to the ones obtained when using the group template, at least with the range of localizers that we used here.

To ensure that our results are robust and not limited to the choice of classifier, we repeated the analysis using a representational similarity analysis (RSA) approach (Kriegeskorte, Mur, & Bandettini, 2008; Nili et al., 2014) and linear discriminant contrasts (LDC). An LDC value between pair of conditions indicates the level of their discriminability, with larger values meaning better discrimination. The difference between the average LDC of all the between-category rule pairs and all the within-category rule pairs (ΔLDC) was calculated per participant and per ROI, for both subject-specific ROIs and group template, with ΔLDC larger than 0 as an indication for rule information. For all localizers as well as when all voxels within the group template were used, the ΔLDC was greater than zero (two tailed: *t_17_* > 3.3, *p* < 0.016, with Bonferroni correction for multiple (4) comparisons), indicating that the distributed patterns of activity conveyed rule information. This was comparable to the trend of distinction type effect seen in the SVM analysis. Importantly, a two-way repeated measures ANOVA with ΔLDC as dependent variable and task (spatial WM, verbal WM, Stroop, all-voxels) and ROI (7) as within-subject factors showed no main effect of task (*F_3, 51_* = 0.3, *p* = 0.7) and no interaction (*F_18, 306_* = 1.1, *p* = 0.35), further supported by individual t-tests of each localizer compared to the group template as well as all possible pairs of localizers (*t_17_* < 0.8, *p* > 0.8, Bonferroni corrected for 6 comparisons). These results indicate that discriminability was similar when using the three localizers to define ROIs in individual subjects and when using all voxels within the group template, in line with the SVM decoding results.

### 3.6 Effect of ROI size on decoding results

In order to examine whether decoding results depend on the number of selected voxels in subject-specific ROIs, as has been observed in the visual system, we performed MVPA for the decoding accuracies across all MD ROIs using different ROI sizes. ROI sizes ranged from 50 to 400, in steps of 50. A three-way repeated measures ANOVA was done with factors: task (spatial WM, verbal WM, Stroop), ROI size (8) and distinction type (2: within- and between-category). There was a main effect of ROI size (*F_7, 119_* = 11.6, *p* < 0.001), no main effect of task (*F_2, 34_* = 0.1, *p* = 0.9) and no interaction between ROI size and task (*F_14, 238_* = 0.7, *p* = 0.7), with the latter indicating that the difference between ROI sizes was the same for the different localizers. We then pooled the data across the three localizers and rule distinctions in order to visualize the main effect of ROI size (**Figure 6**). Classification accuracies tended to be lower for the smaller ROI sizes, and particularly for 50 and 100 voxels, but overall decoding levels were stable with similar decoding accuracies for ROI size of 150 voxels and above. For all ROI sizes, classification accuracy was above chance (*t* > 3.4, *p* < 0.003).

**Figure 6.**
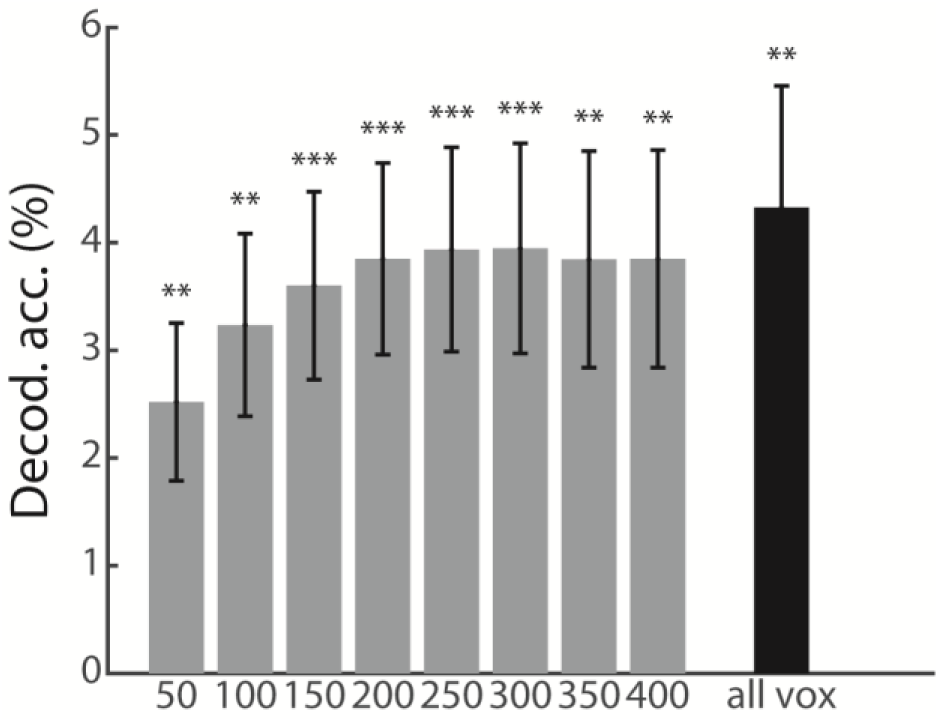
Decoding accuracy for rule distinction for different ROI sizes. Decoding accuracy above chance (50%) is presented for the average of all the localizers, MD ROIs and distinction types. The decoding accuracy level using all the voxels in the MD ROIs (group template) is shown for reference at the rightmost bar. Error bars indicate SEM. ** = p < 0.01, *** = p < 0.001.

## 4. Discussion

In this study we tested for the effect of using individually-defined ROIs within the cognitive control frontoparietal MD network compared to a group template on both univariate and multivariate results in a rule-based criterion task. We systematically tested three localizer tasks (spatial WM, verbal WM, and Stroop) and used a group-constrained subject-specific approach to define ROIs at the single subject level using a conjunction of the independent localizer data and a group mask (Fedorenko et al., 2010). The primary benefit of the proposed approach is the use of both a group template that ensures spatial consistency across participants, as well as individual-subject activation patterns within the group template that provide more focused targeting of MD voxels. We showed a clear benefit for using the subject-specific ROIs compared to when using the group template for mean univariate activity. For multivariate discriminability measures, however, the results were similar for the individually-defined ROIs and for the group template, with no clear benefit (i.e. increased decoding accuracy), or cost (i.e., reduced discriminability), for the subject-specific ROIs approach. Despite differences between the localizers in their spatial activation patterns of the MD network at the individual subject level, we observed similar performance for both univariate and multivariate measures for all three localizers, demonstrating that the choice of localizer did not make a difference to the obtained results. Overall, our results demonstrate that using individually-defined ROIs is a useful way to maintain, or even increase in the case of univariate activity, the sensitivity of broad group templates with an added specificity of activations at the individual subject level.

Our univariate results show that the observed activity is larger when ROIs are defined at the individual level compared to using a group template. These findings replicate and generalize previous findings in other systems such as language and vision, as well as in simulated data (Fedorenko et al., 2010; Nieto-Castañón & Fedorenko, 2012; Saxe et al., 2006). When using the group template, many voxels outside the subject-specific functional signature of the studied region at the individual level are included in the analysis. These voxels average-out the overall activity level, reducing its sensitivity to detect actual changes (Nieto-Castañón & Fedorenko, 2012). Previous studies showed that boundaries between parcellated networks varied across individuals (Glasser et al., 2016; Schaefer et al., 2018; Yeo et al., 2011), further supporting the idea that overall average of activity within a group template may not capture changes that can be observed when ROIs are localized in individual subjects. In addition to the marked increase in mean activity when subject-specific ROIs were used, we also observed larger between-subject variability compared to when the group template was used, with similar variability for all localizers. This difference in variability may be driven simply because averaging is done across many more voxels in the group template, but other factors may include differences between the localizer tasks and the criterion task related to functional specialization within MD regions (Assem, Glasser, Essen, & Duncan, 2019; Crittenden et al., 2016; Dosenbach et al., 2007; Nomura et al., 2010; Shashidhara et al., 2019). Although in our data the increase in mean activity when subject-specific ROIs was detectable and not impaired by the larger variance, future studies may want to take this information into account.

Our main focus in this study was the multivariate results in the MD network and whether individually-localized ROIs will result in a benefit, as seen in pattern discriminability measures, similar to the one observed in the univariate results. Most studies that used MVPA for fMRI data across the frontoparietal network used a group template as ROIs, implemented in a variety of ways. These include, among others, using all voxels within the functionally-defined group-average MD ROIs (Muhle-Karbe, Duncan, De Baene, Mitchell, & Brass, 2017; Woolgar, Hampshire, Thompson, & Duncan, 2011), defining areas of interest based on univariate or multivariate effects of part of the data of the main task and testing on another (Ester, Sprague, & Serences, 2015; Etzel et al., 2016), centering spheres on peak activation loci (Fox, Snyder, Barch, Gusnard, & Raichle, 2005), resting-state networks (Cole et al., 2013), searchlight algorithm (Cole, Ito, & Braver, 2016), and anatomical landmarks in conjunction with group-level univariate contrast (Curtis, Cole, Rao, & D’Esposito, 2005). Because boundaries between specialized areas and networks vary between individuals (Glasser et al., 2016; Schaefer et al., 2018; Yeo et al., 2011), and specifically for the MD network (Fedorenko et al., 2012), the individual ROIs approach has the potential to improve our ability to reliably discriminate patterns of activity within this network. Our results, however, showed that this is not the case. In contrast to the univariate results, pattern discriminability was largely similar when using the individual ROIs and the group template. Our criterion task was chosen to allow for more subtle distinctions between between-category and within-category rule pairs that have been previously observed across the MD network (Crittenden et al. 2016). In the task used by Crittenden et al. (2016), the stimuli were presented at the same time as the colored frames, therefore the between-category and within-category rule pairs were confounded with the category of the stimuli. To avoid this confound, a later study used a variation of the task presenting only colored frames first, and the stimuli later without the cue (Smith et al., 2018). Using this cue and stimulus separation resulted in no difference between the between- and within-category rule pairs, despite strong decoding overall across the MD network. We investigated whether the subject-specific ROIs might show more subtle distinctions between rule pairs. However, the differences between the between-category and within-category rule pairs were similar when using the individual ROIs and the group template, with no benefit for the subject-specific ROIs, for both mean decoding levels and between-subject variability. Importantly, using the individual ROIs for MVPA did not lead to lower discriminability. The preserved accuracy levels demonstrated that the increased localization at the individual subject level did not come at a cost of reduced decodability and maintained the sensitivity to detect task-related neural representations. We note that these results do not depend on the choice of classifier. To ensure the robustness of our results, we used linear discriminant contrast (LDC) in addition to SVM to measure discriminability and differences between localizers and observed similar results. Therefore, the similar multivariate performance when using individual ROIs and the group template was independent of the choice of the MVPA method. Relatedly, rather than overall changes in decoding levels, a potential benefit of using individually-localized ROIs could be an increased likelihood for decoding above chance for individual subjects, driven by reduced within-subject variability of the data. This may be reflected in an increased observed prevalence in the sample, meaning that more subjects show above-chance decoding. Such changes in prevalence can be tested using permutation-based information prevalence inference (Allefeld et al., 2016). However, this analysis is susceptible to small differences in the minimum decoding accuracy in the sample and was not applicable for low decoding accuracies above chance as in our case. Nevertheless, a potential increase in prevalence when individual ROIs are used can be assessed in studies with higher decoding levels. It is noteworthy that despite the preserved decodability when individual ROIs are used, adding a localizer task to the scanning session may have other costs: collecting the additional data in the scanner is expensive, it required extra time of the subjects in the scanner, and it may reduce the amount of data collected for the main task of interest when time in the scanner is limited.

One possible explanation for the similarity of multivariate results obtained for the individual ROIs and the group template is that voxelwise distributed pattern across the entire MD template captured the information related to rule decoding well in the criterion task that we used, with no need for further refinement of the voxels selected for this decoding. Previous studies indeed showed that multivoxel discrimination may also be driven by voxels outside the focused regions of increased univariate activity (Haxby et al., 2001; Kriegeskorte et al., 2006). Another related point is the decoding accuracies that we observed when using the group template, which were at a level similar to what has been shown to be the base rate for the frontoparietal network (Bhandari et al., 2018), therefore potentially limiting our ability to identify increases in decoding levels. Notably, an important limitation of our data is the use of only one criterion task due to limited time in the scanner. It is possible that multivariate benefits or costs of the individual ROIs will be observed for other tasks, and future studies will be required to generalize our results in that respect. We note that since our analysis focused on the fixed-duration cue phase of the trials, our results are well controlled and not driven by behavioral responses such as reaction times. More generally, the underlying factors that contribute to pattern discriminability in the frontoparietal network are not yet well understood (Bhandari et al., 2018), with some previous data showing clear limitations of fMRI decodability compared to what is observed in single-unit data in other brain systems (Dubois, de Berker, & Tsao, 2015). Our data provide another tier of evidence to better understand the relationship between the spatial organization and activity of the MD network at the micro and macro levels and pattern decodability using fMRI.

In this study, we systematically tested three localizers: a spatial WM, verbal WM and a Stroop task. We used tasks that capture a core cognitive aspect associated with the MD network and have the potential to be used as general functional localizers. As expected, and in line with previous results (Fedorenko et al., 2013), all three localizers showed increased activity in the MD network, thus confirming their suitability to serve as localizer tasks. Activation patterns of the localizers in individual subjects were highly reliable, as reflected in high correlations across voxels between the two runs of each localizer. In contrast, these correlations were substantially reduced across subjects, as is also shown in the whole-brain overlay maps, demonstrating the need for a subject-specific ROI definition approach. In line with previous data (Fedorenko et al., 2013), there were substantial correlations between activation patterns of the different localizers (computed within each subject), but these were lower than those between runs of the same localizer. The Stroop task evoked a weaker, and less MD-focused pattern of activity, and had lower reliability between runs. These differences compared to the other two localizers were small and in some cases only marginally significant, but may imply that the Stroop task captured the MD network less well. The spatial differences in recruitment between the localizers may reflect differential functional preferences for cognitive demands and constructs across MD regions (Assem et al., 2019; Crittenden et al., 2016; Dosenbach et al., 2007; Nomura et al., 2010; Shashidhara et al., 2019). Specifically, the two working memory tasks might reflect a more similar cognitive construct compared to the Stroop task that involves conflict monitoring and inhibition. Another possible explanation for the difference could be related to the difficulty manipulation in the tasks. While increase in difficulty level in the working memory tasks was simply controlled by increasing the number of highlighted cells in the grid or numbers, this manipulation in the Stroop task was operationally less well defined.

An important aspect of the individual ROI approach that we used is the ability to control for the ROI size. It has been previously shown that increased ROI size leads to increased classification levels in the visual system, highlighting the need to control for ROI size when comparing results across ROIs (Eger et al., 2008; Erez & Yovel, 2014; Said et al., 2010; Walther et al., 2009). It was not clear whether this is the case for MD regions, which are different from visual regions in multiple respects, and whether more generally the choice of this parameter affects decoding levels. In our data (Figure 6), we observed slightly lower levels of classification for the smaller ROI sizes (50 and 100 voxels), but these stabilized for ROI sizes of 150 voxels or more, in line with previous reports (Erez & Duncan, 2015; Shashidhara & Erez, 2019). This does not necessarily mean that such high dimensionality is required to reach maximal decodability without over-fitting, an issue that other studies have looked into more formally (Ahlheim & Love, 2018). Importantly, controlling for ROI size may be essential when comparing MD regions to each other as well as to other brain systems, such as visual areas, further emphasizing the importance of using a method that enables such control.

Based on our systematic comparison of the localizer tasks and the effect of using individual ROIs for univariate and multivariate analysis, several recommendations can be drawn for future studies of the MD network to inform researchers in the field while designing a study:

1. Using a localizer task and subject-specific ROIs may be highly beneficial when the question of interest concerns univariate activity and level of recruitment of the MD network. For studies that are designed to use both univariate and multivariate approaches, subject-specific ROIs can be used to benefit the univariate results without impairing multivariate discriminability.
2. If only MVPA is used, using subject-specific ROIs may not improve discriminability. Nevertheless, researchers might still want to use subject-specific ROIs for multivariate analysis when recruitment of the MD network at the individual subject level is important for theoretical reasons (e.g. separating MD activity from adjacent specialized areas such as the language system). Otherwise, localizer data may not be required, thus saving scanning costs and time in the scanner.
3. Localizer data and individual ROIs may be important in studies of the MD network where control for ROI size is required, even if only multivariate analysis is used. This could be the case, for example, when comparing results to other brain systems where decoding levels are sensitive to the ROI size, such as the visual system.
4. Choice of localizer: although the overall results were similar for the three localizer tasks, our data shows that the Stroop task captured the MD network less well than the two other localizer tasks. Therefore, either the spatial or the verbal working memory tasks may be more suitable as candidate localizer tasks to identify individual MD regions.
5. ROI size: since decoding accuracies in our data were stable for ROI size of 150 voxels and above, any choice or ROI size of 150 or larger may provide similar multivariate results.
6. The amount of localizer data that should be collected is another factor that needs to be considered. Within-subject patterns of activity of the two localizer runs were highly similar for both the spatial and verbal working memory tasks (reliabilities of > 0.7), but not identical. These correlations imply that using two runs of the localizer task may be required to capture MD activity in individual subjects, but one run may be sufficient in cases where time in the scanner is a major constraint. We note that statistical comparison of the results when using two runs or one run only to define individual ROIs requires three runs for each localizer to avoid dependencies in the data, and therefore could not be done in our study. This question may be addressed more formally in future studies.

Beyond the localizer tasks that we used here, different variations of the individual ROI approach can be designed for future studies, offering a balance between the need of a consistent definition of regions across participants, and perhaps studies, and the localization at the individual participant level. Such variations can be designed depending on the research question, and can be used with both univariate and multivariate analyses while avoiding double-dipping (Kriegeskorte, Simmons, Bellgowan, & Baker, 2009). For example, data from two or more localizer tasks can be combined, as we demonstrated here. Combining data across localizers could lead to capturing core parts of the MD system, thus reflecting the multiple-demand nature of the selected voxels (Assem et al., 2019; Duncan, 2010, 2013; Fedorenko et al., 2013). An even more cognitively diverse variation can be a localizer that consists of multiple tasks within the same run with a similar manipulation of difficulty level. On the other end of this scale, a localizer task can be designed to target a specific cognitive aspect of interest, and constraining activation patterns by a group template will ensure that the areas of interest are within the boundaries of the MD network.

In summary, we used three independent localizer tasks to define subject-specific ROIs and test the effect of using the individual ROIs compared to a group template on univariate and multivariate effects in a rule-based criterion task. The univariate results in the criterion task greatly benefitted from using individual ROIs compared to using the group template. In contrast, multivoxel task-related representations as measured with MVPA were similar for the individual ROIs and the group template, and for all localizers. There was no benefit of increased pattern discriminability, as well as no cost of reduced discriminability, for the individually-localized ROIs. The group-constrained individually-defined ROIs offer a refined and targeted localization of each participant’s MD regions based on the individual’s unique functional pattern of activity, while ensuring that similar brain regions are studied in all participants. Pushing forward towards standardization in the field, our study provides important empirical evidence for researchers using both univariate and multivariate analysis of fMRI data to study the functional organization of the MD network.

## Acknowledgements

This work was funded by a Royal Society Dorothy Hodgkin Research Fellowship (UK) to Yaara Erez (DH130100). Sneha Shashidhara was supported by a scholarship from the Gates Cambridge Trust, Cambridge, UK. Floortje Spronkers was supported by an Erasmus+ Traineeship grant and a Stichting A.S.C. Academy grant. We thank John Duncan and Daniel Mitchell for fruitful discussions and advice throughout the study.

The authors declare no competing financial interests.

